# Integrated genomic analysis of NF1-associated peripheral nerve sheath tumors: an updated biorepository dataset

**DOI:** 10.1101/2024.01.23.576977

**Authors:** Jineta Banerjee, Yang Lyu, Stavriani C. Makri, Alexandra J. Scott, Lindy Zhang, Ana Calizo, Kai Pollard, Kuangying Yang, John M. Gross, Jiawan Wang, Adam S. Levin, Allan J. Belzberg, Carlos G. Romo, Robert J. Allaway, Jaishri O. Blakeley, Angela C. Hirbe, Christine A. Pratilas

**Author notes:** These authors contributed equally (Jineta Banerjee, Yang Lyu, Stavriani C. Makri).

## Abstract

Neurofibromatosis type 1 (NF1) is an inherited neurocutaneous condition that predisposes to the development of peripheral nerve sheath tumors (PNST) including cutaneous neurofibromas (CNF), plexiform neurofibromas (PNF), atypical neurofibromatous neoplasms with unknown biological potential (ANNUBP), and malignant peripheral nerve sheath tumors (MPNST). The successful advancement of therapeutic development for NF1-associated PNST necessitates the systematic acquisition and analysis of human tumor specimens and their corresponding model systems. RNA sequencing (RNAseq) and whole exome sequencing (WES) data were generated from 73 and 114 primary human tumor samples, respectively. These pre-processed data, standardized for immediate computational analysis, are accessible through the NF Data Portal, allowing immediate interrogation. This analysis combines new and previously released samples, offering a comprehensive view of the entire cohort sequenced. As a dedicated effort to systematically bank tumor samples from people with NF1, in collaboration with molecular geneticists and computational biologists to advance understanding of NF1 biology, the Johns Hopkins NF1 biospecimen repository offers access to samples and genomic data to promote advancement of NF1-related therapies.

## Background & Summary

Neurofibromatosis type 1 (NF1) is an inherited neurocutaneous condition that predisposes to the development of peripheral nerve sheath tumors (PNST) including cutaneous neurofibromas (CNF), plexiform neurofibromas (PNF), atypical neurofibromatous neoplasms with unknown biological potential (ANNUBP) (1), and malignant peripheral nerve sheath tumors (MPNST). Historically, therapeutic progress for PNF and MPNST has been limited in part due to restricted availability of primary tissues from patients with NF1. The successful advancement of therapeutic development for NF1-associated PNST necessitates ongoing efforts in the systematic acquisition and analysis of human tumor specimens and their corresponding model systems.

The Johns Hopkins NF1 biospecimen repository strives to address this gap through the collection of diverse biospecimens from patients with NF1. Patients with clinically or genetically confirmed NF1 having a clinically indicated surgical resection or biopsy of any NF1-associated tumor were invited to participate in an institutional review board (IRB) approved study for the collection and sharing of tissues and specimens. Tumors are routinely collected, assessed by a study pathologist, and banked in the laboratory as flash frozen tissues, paraffin embedded blocks or slides, DNA and RNA, or single cell suspensions. Efforts are ongoing to create cell lines and patient derived xenografts (PDX) from primary human tissues (2–4). Clinical data for participating patients are fully annotated in a database that corresponds to banked tissue specimens. Applications for access to biospecimens, genomic data, and disease models, as well as de-identified clinical and molecular data are reviewed and approved with IRB oversight, to allow internal and external sharing to promote research collaboration.

Our comprehensive efforts to generate a genomically and clinically annotated NF1 biospecimen repository, for the purpose of sharing resources, to promote collaborative research, and make specimens and genomic data available to the NF1 research community, were previously reported (5). As of June 2024, the Johns Hopkins NF1 biospecimen repository now comprises a total of 390 specimens including PNF (n=91), ANNUBP (n=7), MPNST (n=66), CNF (n=118), diffuse neurofibroma (diffuse NF, n=44) or another NF1-associated tumor (n=64). A full specimen inventory is available online and investigators can request specimens by providing a statement of scientific rationale and intended use (data use statements: https://doi.org/10.7303/syn4939902). More than 70 requests for sharing of available specimens, models, and/ or genomic data have been fulfilled to date, supporting the existence of this biobank as a successful, active, and widely recognized resource for promoting a diverse set of research needs in the NF1 community.

The dataset presented herein includes whole exome and RNA sequencing, with a validation analysis meant to provide the scientific community with an overview of available samples and their potential utility. Our validation of the data confirms findings that are concordant with available literature regarding the genomics of NF1-associated tumors and validates the biospecimen repository genomic data as a useful resource for further downstream analysis by NF1 researchers.

## Methods

### Tumor sample collection and sequencing

#### Patient enrollment

All research was conducted according to a Johns Hopkins Hospital (JHH) institutional review board (IRB)-approved protocol, IRB00096544.

Patients with clinically or genetically confirmed NF1 having a clinically indicated surgical resection or biopsy of any NF1-associated tumor (including, but not limited to, CNF, PNF, ANNUBP, MPNST, other NF1-associated neoplasm) were invited to participate. Written informed consent was obtained from all patients. The JHH IRB-approved consent form includes a statement about the voluntary nature of the research, a description of corresponding clinical data that will be collected for future analysis, and a statement that samples may be shared beyond JHH. Blood was collected from most patients on the day of surgery.

All study subjects, including those without germline genetic testing, who donated samples to the Johns Hopkins NF1 Biospecimen Repository and have been analyzed in the current manuscript have met the NIH clinical criteria for diagnosis of NF1 (6). Out of 54 unique participants listed on Table 1, 16 patients have had blood genetic characterization of their NF1 variant. Out of 16 patients, 3 were diagnosed with NF1 microdeletion syndrome, 3 were found to have 1.4 megabase deletion of NF1 gene, 5 were found to have known pathogenic variants, 2 were found to have NF1 variants that are likely pathogenic, 2 were found to have an NF1 variant of uncertain significance, and 1 had a negative result but the NF1 diagnosis was established based on this patient meeting the clinical criteria.

**Table 1.**
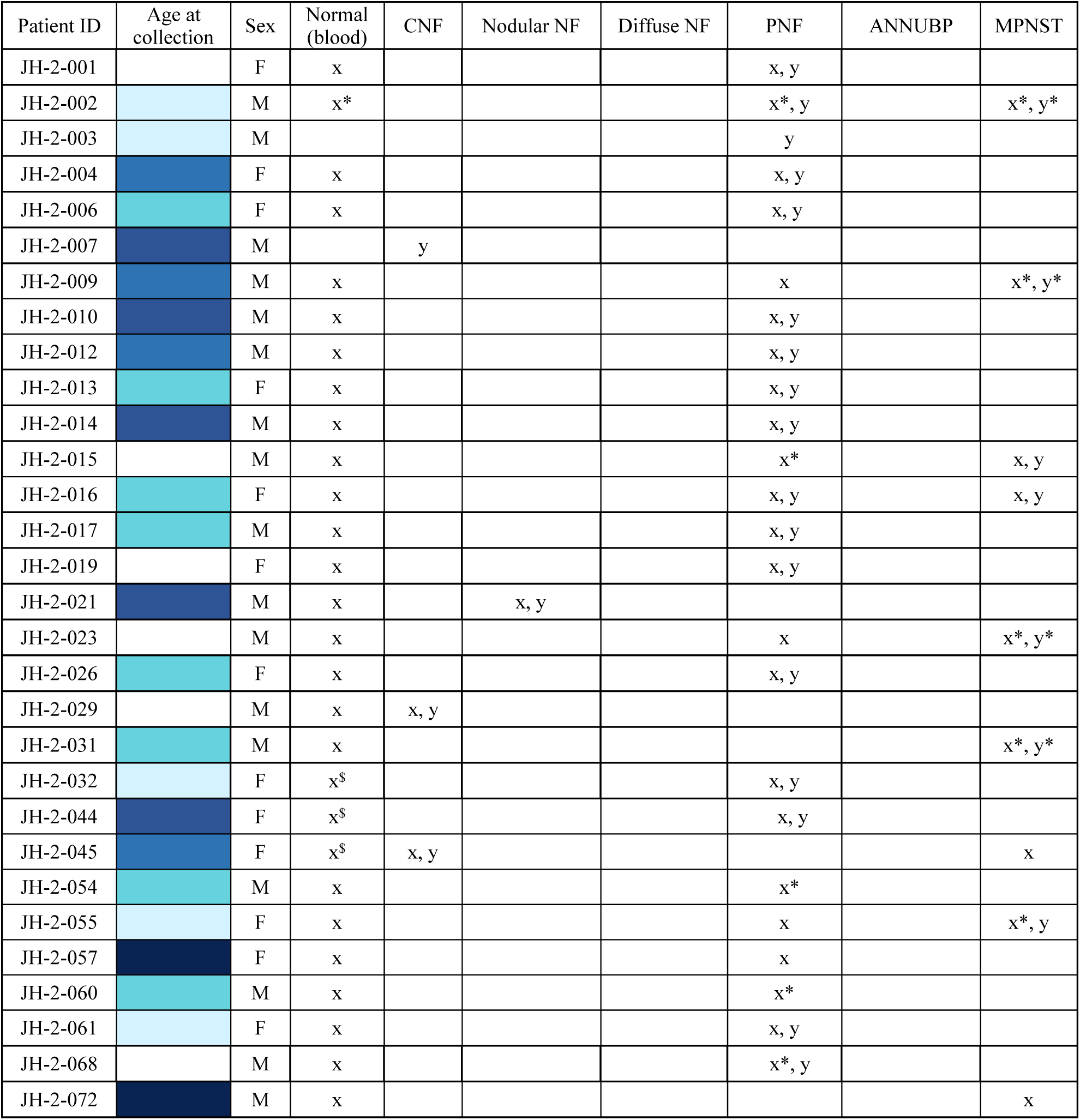
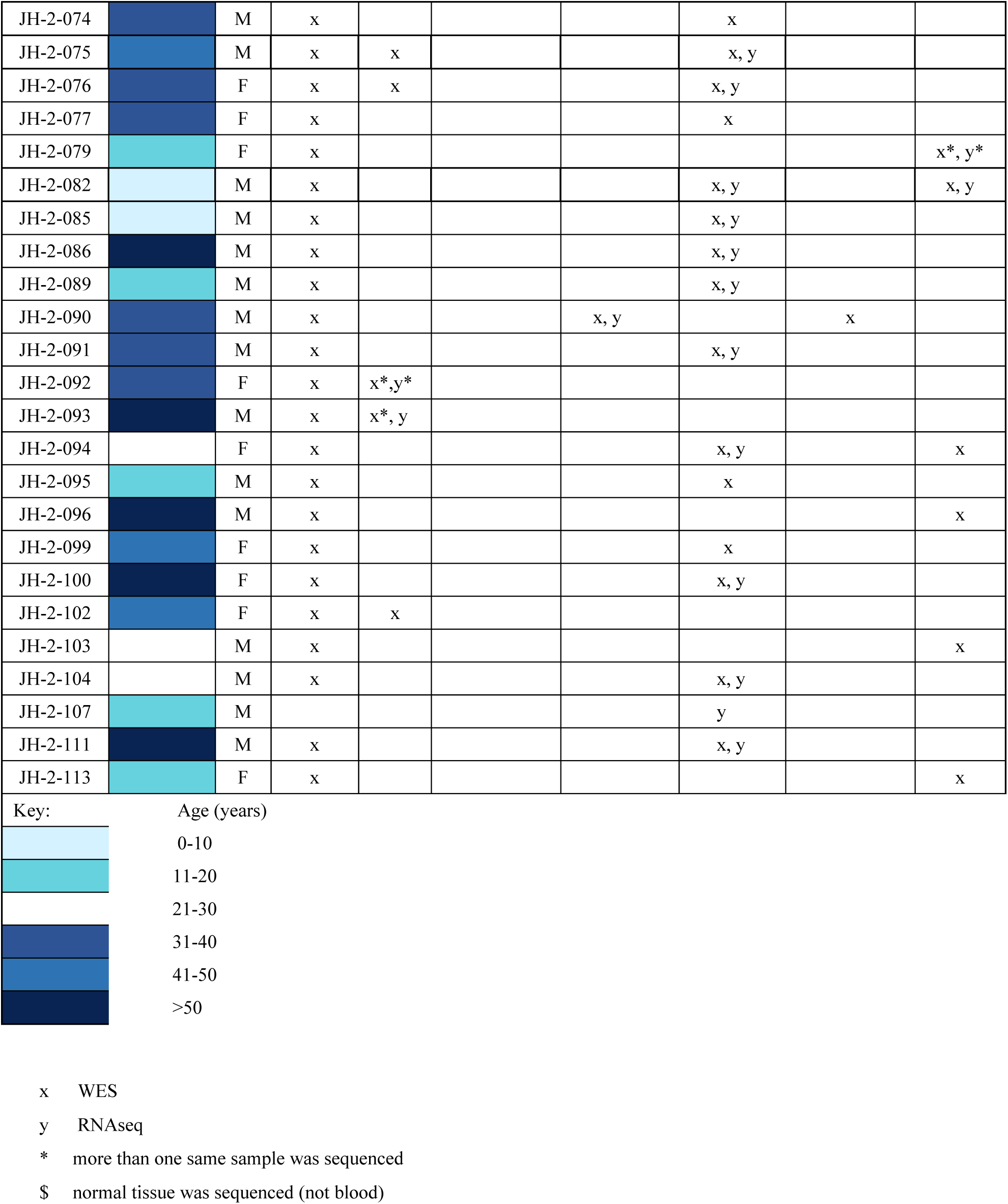
List of samples with RNAseq and/ or WES, including age (in years by range), sex, and type of NF1-associated PNST, included in the current analysis of samples and for which sequencing data are available (inclusive of samples released with Pollard et al, *Sci Data*, 2020 (5)). WES and RNA sequencing are indicated by an “x” and “y” symbol, respectively.

Medical records were retrospectively reviewed for demographic information and information related to the NF1 clinical condition (genetic diagnosis, family history, age of diagnosis), characteristics (phenotypic findings, symptoms), and tumor burden (number and size of NF1-associated tumors). These clinical data are stored in a password protected and de-identified database at JHH.

#### Tumor preservation and quality control

Surgical specimens were couriered to surgical pathology immediately after resection. The study pathologist (JMG) performed immediate inspection of the tumor to ensure that adequate tumor existed for clinical diagnostic needs. Upon approval, tumor pieces were sampled for banking and transported to the research laboratory in harvest medium (RPMI with 20% fetal bovine serum, supplemented with 1% penicillin-streptomycin and glutamine).

Specimens were sized into 5-10mm aliquots under sterile conditions in a biosafety cabinet. Individual aliquots were placed into 10% neutral buffered formalin, cell freezing media (Sigma #C6295), RNAlater (Qiagen #76104), and/ or placed into an empty vial and snap frozen on dry ice. Formalin-fixed paraffin-embedded tissues were stained with hematoxylin and eosin (H&E). H&E slides from each specimen were reviewed by the study neuropathologist to confirm an accurate diagnosis of each preserved specimen.

#### Tumor sample preparation for DNA sequencing

Genomic DNA from blood and tumor was extracted using the QIAGEN DNeasy Blood & Tissue kit. DNA library was constructed using KAPA HyperPrep Kits for NGS DNA Library Prep (Roche #7962363001). Exomes were captured by IDT xGen Exome Hyb Panel v1.0 (IDT #1056115) and the libraries were sequenced by NovaSeq6000 S4 with ∼100X coverage for normal samples and tumor samples.

#### Tumor sample preparation for RNA sequencing

RNA was isolated from tumor by using TRIZOL. Samples containing at least 100ng total RNA with RNA Integrity Number (RIN) > 6.5 were sequenced. For RNA extraction, the first batch of samples were prepared with MGI/GTAC RiboErase method. Depletion was performed with KAPA RiboErase HMR (Roche #07962274001). Library preparation was performed with a WUSTL in-house preparation protocol. For the second batch, samples were prepared with Illumina TruSeq Stranded Total RNA Library Prep Gold kit (#20020599) which includes RiboZero Gold depletion. Library preparation was performed according to the manufacturer’s protocol. Fragments from both batches were sequenced on an Illumina NovaSeq-6000 using paired-end reads extending 150 bases.

### Whole exome sequencing data processing for somatic variant calling and analysis

Raw fastq data files were quality checked using FastQC v0.11.9 and a report was generated using MultiQC v1.8. Fastq files were aligned to GRCh38 using BWA 0.7.17-r1188 (7). Duplicates were marked using GATK MarkDuplicates, and bases recalibrated using GATK BaseRecalibrator and GATK ApplyBQSR (GATK v4.3.0.0) (8–10). Somatic single nucleotide variants (SNVs) were then called using Strelka2 software (Strelka v2.9.10) (11) and Mutect2 (GATK v4.4.0.0) (12). Strelka 2 was shown to have high precision and recall for SNVs compared to others as tested in the precision FDA challenge (11, 13). The variants were annotated using Variant Effect Predictor (VEP v99.2) and converted to mutation annotation format (MAF) files using vcf2maf (vcf2maf v1.6.21). All of these steps were completed on Nextflow Tower running the standardized nf-core pipeline sarek v3.1.2 (14).

### Variant calling from samples with paired normal in a separate batch

The samples contained in syn52659564 and **Supplementary Table 1** were special cases where the tumor samples and the paired normal samples were sequenced in two different batches with different sequencing library kits and thus had two different browser extensible data (BED) files for target capture. For the above samples, first the BED file from JH_batch1 was lifted over from hg19 to hg38 coordinates using the UCSC liftOver tool (15) and the liftover chain file named ‘hg19ToHg38.over.chain.gz’ and then sorted using the sort function from bedtools suite. Then a common region BED file was generated using the intersect function in bedtools suite with at least 50% overlap in intervals between the BED file from JH_batch1 and WU_batch1. The common region BED file (syn52594249) was then used to call somatic variants from these tumor-normal pairs using the sarek v3.2.2 pipeline and Strelka2 and Mutect2 variant callers.

### Consensus variant calling using SomaticSeq

To avoid detection of false positive variant calls in the samples, we identified and reported the variant calls that had consensus between Strelka2 and Mutect2 callers. To identify these consensus calls we used a previously benchmarked method called SomaticSeq (16). Once consensus calls for SNVs and insertions and deletions (indels) were identified, the variants in respective variant call formats (VCFs), were annotated using vcf2maf in the sarek v3.2.2 pipeline to generate a merged MAF file that was used for further analysis. The consensus variants were filtered for high confidence calls (PASS filtered) with population allele frequency below 0.05 to identify robust variant calls excluding common variants.

### RNAseq data processing and analysis

Raw fastq files were processed using nf-core/rnaseq (v3.11.2) (17) and quantified using salmon: v1.10.1 (18). ComBat from sva R package (v3.42.0) was used for batch correction (19). DESeq2 (v1.34.0) was applied to call the differentially expressed genes (|fold change| > 1.2, adjust *P* < 0.05), and ggplot2 R package (v3.3.6) was applied to draw volcano plots. Principal component analysis (PCA) and uniform manifold approximation and projection (UMAP) (20) were applied for the visualization of samples. Gene set enrichment analysis (GSEA) was conducted by using R-package fgsea (v1.20.0) [13]. ggplot2 (v3.3.6) was used for the gene volcano plot and pathway enrichment plot [14]. Pheatmap (v1.0.12) was applied to plot gene expression heatmap. Singscore (v1.14.0) was applied to calculate single sample gene signature score (21).

### Evaluation of relatedness in genomic and transcriptomic data

Somalier (v0.2.17) (22) was used to evaluate the relatedness between WES and RNAseq samples. The *extract* function was used to identify informative sites from the WES cram files and RNAseq bam files, and then the *relate* function was used to calculate statistics on the number of shared genotypes at informative sites and the coefficient of relatedness between all pairs of samples. One WES sample, JH-2-009-2578C-A, did not contain data at a sufficient number of polymorphic sites to calculate relatedness to other samples and was excluded from further analysis. The pheatmap R package (v1.0.01) was used to generate heatmaps of the relatedness comparing each primary human WES sample exactly once to each primary human RNAseq sample and the ggplot2 package was used to generate scatter plots. Cell line and PDX samples were excluded from the heatmap.

#### Data Records

The goal of this project is to bank blood and tumor samples with accompanying clinical data from patients with NF1-associated neoplasms. These tissues, and accompanying genomic data, represent a publicly available resource shared routinely with researchers, to promote ongoing collaborative science for therapeutic discovery. A total of 187 samples (plus 71 corresponding normal samples) from banked tissues have been sequenced, including whole exome on a total of 114 tumors (33 MPNST, 4 ANNUBP, 57 PNF, 9 diffuse NF, and 11 CNF) and RNA sequencing on a total of 73 tumors (18 MPNST, 1 ANNUBP, 38 PNF, 9 diffuse NF, 6 CNF, nodular NF 1). These genomic data are now publicly available to NF1 researchers to support an array of important lines of scientific inquiry.

With this publication, we are releasing curated datasets of pre-processed WES and RNAseq data that are ready for general re-use. These datasets are processed using standardized publicly vetted processing workflows to reduce the efforts of data re-use. Multiple variant callers and well known RNAseq quantification methods have been used to generate datasets designed for maximum reusability. Additionally, these datasets are well annotated with clinical metadata to enable appropriate re-analysis of the data as needed.

Included in this validation dataset are primary tumor samples sequenced using WES (82 total: MPNST 24; ANNUBP 1; PNF 45; diffuse NF 1; CNF 10, nodular NF 1) and RNAseq (51 total: MPNST 14; PNF 29; diffuse NF 1; CNF 6, nodular NF 1). All existing RNAseq and WES raw data for these tumors, as well as 4 MPNST cell lines and 2 PDX, have been deposited in the NF Data Portal:

https://nf.synapse.org/Explore/Studies/DetailsPage/Details?studyId=syn4939902.

All the raw data files are available on Synapse.org

(WES: dataset format: https://doi.org/10.7303/syn53132831.1,

RNAseq: dataset format: https://doi.org/10.7303/syn53133024.1).

All processed data visualized in this article are available on Synapse.org: (WES: Somatic Variants Mutect2: https://doi.org/10.7303/syn53149144.2,

Somatic Variants Strelka2: https://doi.org/10.7303/syn53149128.1,

RNAseq quantification files: https://doi.org/10.7303/syn53140231.1,

RNAseq counts files: https://doi.org/10.7303/syn53141534.1).

All curated datasets can be viewed and accessed through the NF Data Portal: https://bit.ly/nfbiobankdatasets.

#### Technical Validation

##### Patients selected for tissue banking satisfied NF1 clinical diagnostic criteria

Participating patients either 1) met the clinical criteria for neurofibromatosis type 1 AND had genetic confirmation of NF1 gene alteration, or 2) met the clinical criteria for NF1 only, without clinical genetic testing. Clinical and demographic information on the patients whose tumors have been genomically characterized, including sex and age by range, are included in **Table 1**. Specific ages at the time of tumor collection are not provided to protect patient identities given the rarity of some NF1-associated conditions. Sample and file information can be found in Synapse.org datasets (WES: syn53132831.1, RNAseq: syn53133024.1; for more details, refer to the Data Records section) and accessed on the NF Data Portal for further analysis.

Relatedness of WES and RNAseq data from individual patients was confirmed as a measure of quality control.

As a quality and validation step, we used Somalier (22) to calculate the coefficient of relationship (“relatedness”) between WES and RNAseq samples, comparing each WES sample exactly once to each RNAseq sample in order to detect sample swaps. We found that all sample comparisons that originated from different individuals had a relatedness coefficient <0.9. Based on this and the clinical annotations of the samples, we set a threshold of relatedness coefficient ≥0.9 to be the confirmation that a pair of WES and RNAseq samples were indeed from the same individual (**Figure 1A**).

**Figure 1.**
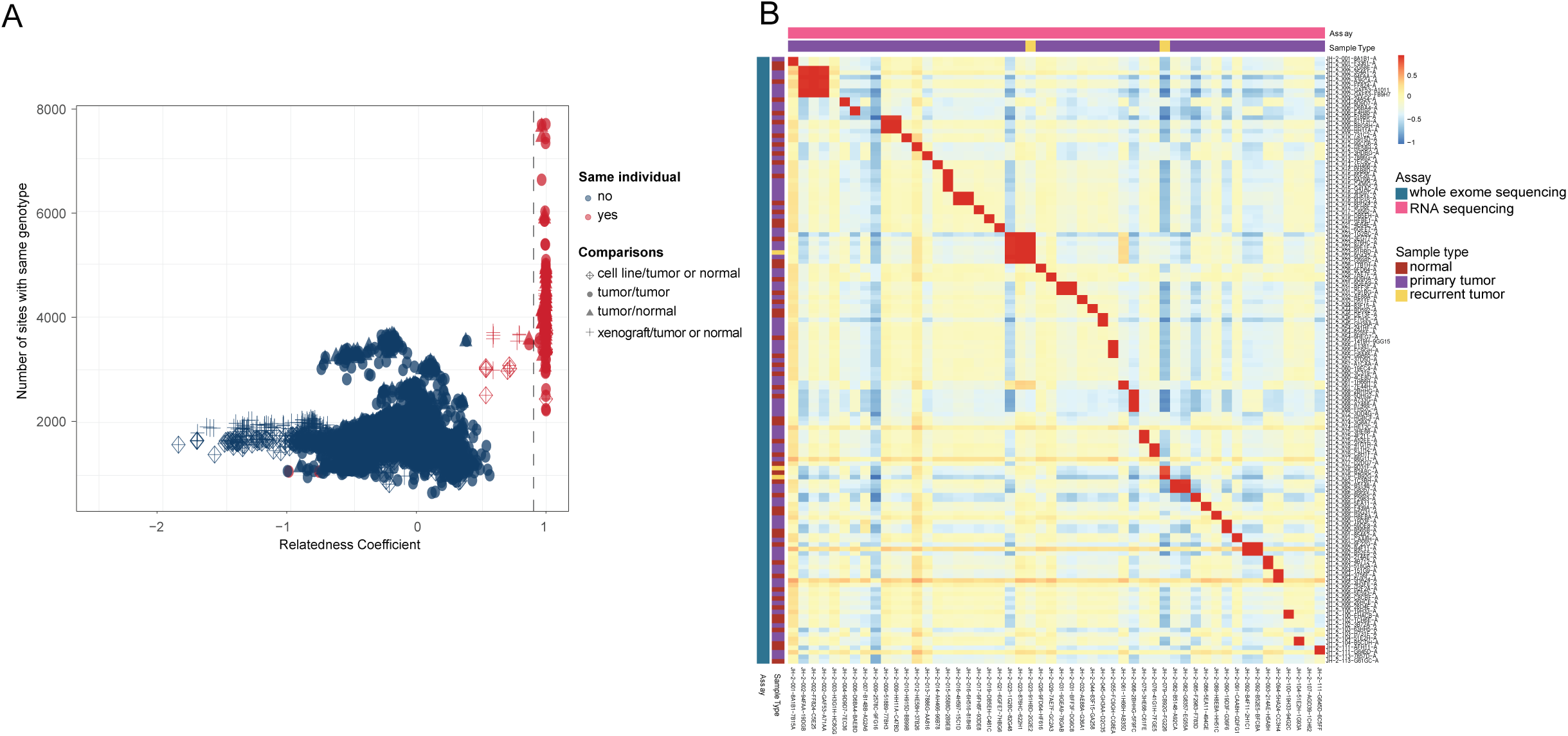
Relatedness between WES and RNAseq samples. A) A scatter plot of the relatedness coefficient versus the number of sites with the same genotype between two samples, as calculated by Somalier. Comparisons between two samples annotated as being from the same individual are shown in red and those from different individuals are colored blue. The shape of the point indicates the tissue of origin of the samples compared. The vertical line indicates the cutoff value (≥ 0.9) used to determine whether two samples are from the same individual based on the relatedness coefficient. B) A heatmap of the relatedness coefficient of pairwise comparisons between WES and RNAseq data from primary human samples as shown in (A), excluding cell lines or xenografts derived from these samples. RNAseq samples are shown on the x-axis and WES samples are on the y-axis. The relatedness coefficient is displayed according to the color scale bar shown in the figure.

With this threshold in place, all but three corresponding samples from the same patient were found to meet the definition of relatedness (**Figure 1A,B**). One RNAseq sample from a recurrent tumor, JH-2-079-CB92G-FG226, had a relatedness coefficient of 0.86 to both the normal WES sample from the same individual and to a WES sample from a different tumor in the same individual. However, it had high relatedness (0.997) to the WES sample generated from the same recurrent tumor. Two additional RNAseq samples that originated from one individual had lower than expected relatedness to other samples from that individual. On examination of the clinical annotations, we found that JH-2-002-GAF53-2E57B was an RNAseq sample of a patient-derived cell line and JH-2-002-GAF53-3BAGC was an RNAseq sample of a xenograft developed from the tumor sample. This distinction suggested that the threshold of relatedness coefficient determined by our analysis was sensitive enough to detect differences between primary samples and the cell lines and xenografts derived from them. The above results confirmed that both RNAseq and WES data obtained from the same primary tumor and labeled with the same patient ID were indeed comparable.

##### Unbiased exploration of somatic variants identifies NF1 among the top genes with variants using WES

WES data from the human tumor samples were haplotype matched to their respective normal samples (**Supplementary Table 2**). To validate the variant landscape of the human tumor samples from the biobank, we consulted six highly cited reports (5, 23–27) to generate a list of 60 priority genes that are deemed to be of key biological importance in NF1-MPNST. Given the differences in detecting limits of variant caller algorithms, we used multiple variant callers to identify single nucleotide polymorphisms (SNPs) and short indels for these genes of interest in the tumor samples. Variants detected using Strelka2 (**Figure 2A,D,E)** exceeded those detected using Mutect2 (**Figure 2B,F,G**). The median number of variants found in the samples using Strelka2 and Mutect2, respectively, are shown in **Figure 2D** (median =129) and **2F** (median=11) and highlight that the majority of the variants detected were missense variants. The top ten genes with variants detected using Strelka2 and Mutect2 callers are shown as **Figures 2E,G**. To find variants that were identified regardless of the variant caller used, we identified consensus variant calls with SomaticSeq – (**Figure 2C,H,I)**. Samples in which no high-confidence variants were detected using at least one variant caller were excluded from further analysis, and those that had consensus calls for variants using SomaticSeq are visualized in **Figure 2C**. A median of 6 variants including missense, nonsense, splice-site, nonstop, and translation start-site variants were found per sample when only consensus calls were considered (**Figure 2H)**. In the consensus calls, *NF1 and SUZ12* genes were found within the top 10 genes with variants with moderate or high impact on gene function among the samples (**Figure 2I**).

**Figure 2.**
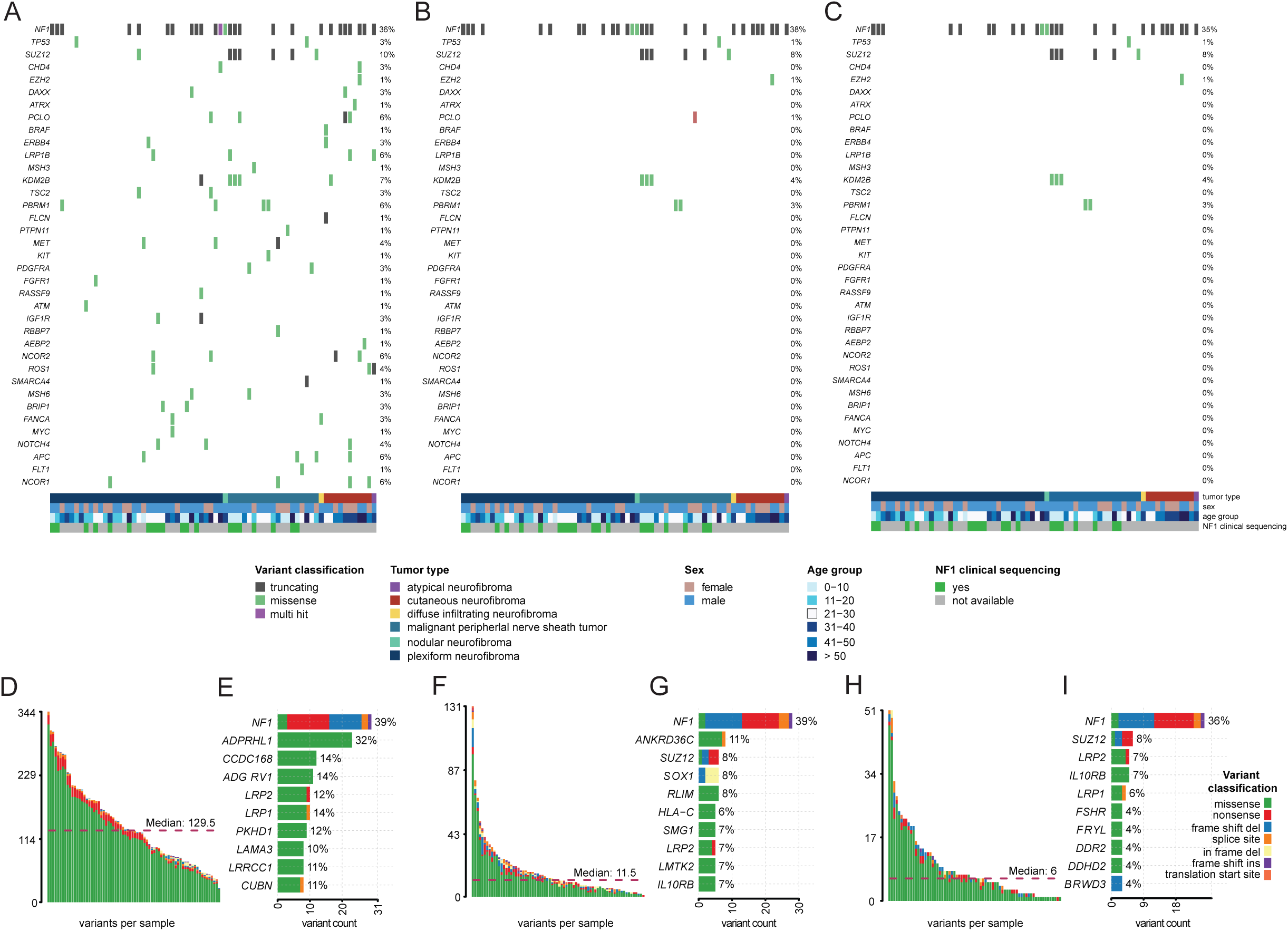
Summary of somatic variants detected in PNST (10 CNF, 1 nodular NF, 1 diffuse NF, 36 PNF, 1 ANNUBP, and 19 MPNST). A) Oncoplot of variants in selected genes of interest (see methods) in the cohort of biobank patients detected using Strelka2. B) Oncoplot of variants in selected genes of interest in the cohort of biobank patients detected using Mutect2. C) Oncoplot of the consensus of variants in selected genes of interest in the cohort of biobank patients detected using SomaticSeq. The status of *NF1* functional inactivation in the patients was determined using either clinical genetic testing or clinical diagnosis and is provided using colored bars in the bottom panel of each oncoplot. (D, F, H) Barplot showing the number of variants per sample, by SNV class (missense, nonsense, splice site, nonstop or translation start site). (E, G, I) Bar plot of the top 10 genes with variants of moderate or high impact in the cohort.

We identified somatic SNVs in the *NF1* gene in 31% of PNF samples and 26% of MPNST samples examined. All consensus variants identified in the NF1 gene were predicted to have moderate or high impact as determined by the Ensembl variant effect predictor (VEP) (28). This suggests that these variants either decreased the protein function without disrupting the protein structure (moderate impact) or disrupted the protein activity through truncation, loss of function, or nonsense mediated decay (high impact) respectively. The genetic loci of *NF1* variants identified in our samples are shown schematically alongside a protein structure representation of the *NF1* gene (**Figure 3A**, PNF shown at top, MPNST shown at bottom). **Figure 3B** represents a diagrammatic rendition of known functional domains in the NF1 protein for illustration of the position of the variants identified relative to the domains of the protein sequence. **Figure 3B** is modified from previously published literature for illustration purposes only (29). We note that variants in the *NF1* gene were detected in 35% of the samples rather than the expected 100% of samples, given their origin from people with the NF1 clinical condition. This detection rate is consistent with prior reports in which *NF1* alterations are identified in 40% of tumors analyzed (30, 31). The inability to detect *NF1* variants in all samples despite a clinical diagnosis highlights the current limitations with WES technology and the many ways in which the *NF1* gene can be altered resulting in the variable clinical presentation of NF1.

**Figure 3.**
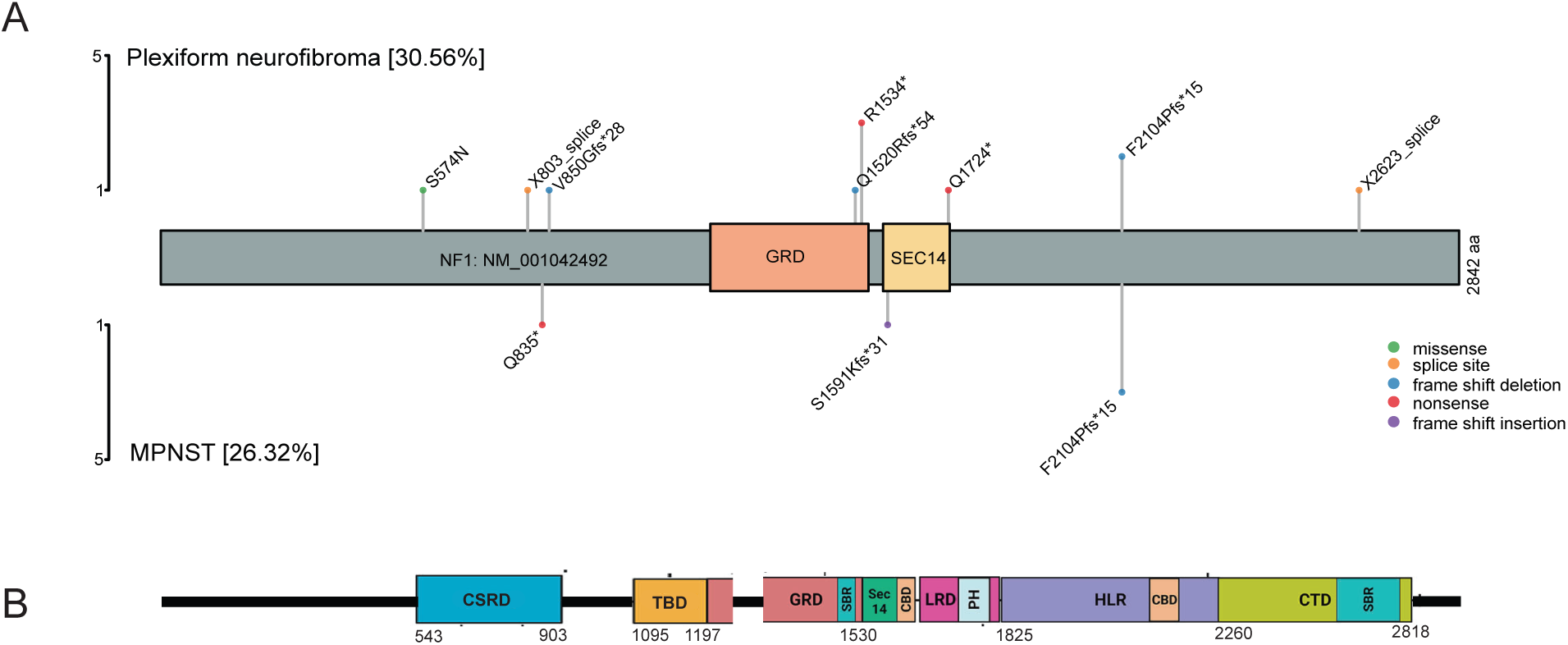
Somatic variants in plexiform neurofibromas and malignant peripheral nerve sheath tumors. A) Lollipop plot showing different positions and protein consequences of variants in the *NF1* gene as detected in this cohort using WES. B) Schematic representation of neurofibromin protein with various functional domains. Domains: CBD caveolin-binding domains, CSRD cysteine and serine rich domain, CTD C-terminal domain, GRD RAS-GAP-related domain, HLR HEAT-like repeats, LRD leucine-rich domain, PH pleckstrin homology, SBR syndecan-binding region, TBD tubulin-binding domain (adapted from Mo et al, 2022, with permission) (29).

##### MPNSTs demonstrate enrichment in pathways associated with progression from plexiform neurofibroma to malignancy

We employed PCA to cluster the samples based on the gene expression data (**Figure 4A**). The majority of CNF tumors clustered together, distant from the MPNST samples, indicating divergent gene expression profiles between the two tumor types. Most PNF and MPNST samples clustered nearby each other, possibly indicating a range of biological behavior in PNF, which are known to have a propensity for malignant progression in some patients, or possibly MPNST with less aggressive biology or potential for metastasis. Interrogation of the clinical outcomes for specific patients from whom these PNF were obtained would take years of follow up and is therefore outside the scope of this analysis.

**Figure 4.**
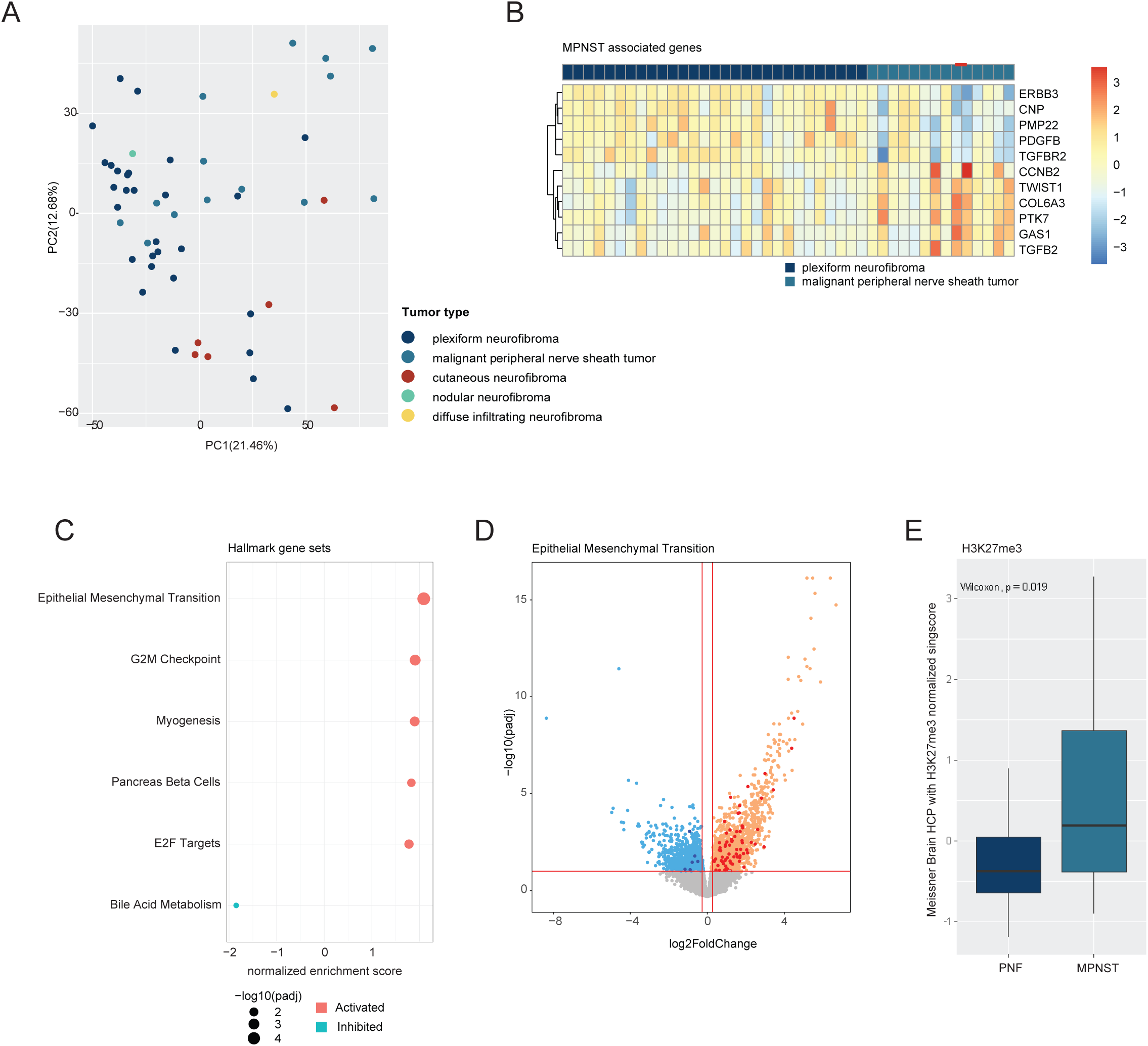
A) Principal component analysis (PCA) plot illustrating the distribution of all tumors based on RNAseq gene counts. B) Heatmap displaying the expression patterns of differentially expressed genes associated with MPNST in comparison to PNF. C) Dot plot showing significantly enriched pathways in Hallmark gene sets in MPNST vs. PNF (adjusted *P* < 0.01). D) Volcano plot for the most significant Hallmark pathway (epithelial-mesenchymal transition) highlighting differentially expressed genes in MPNST vs. PNF (adjusted *P* < 0.01). E) Boxplot of the single sample gene signature score (singscore) in one significant pathway in MPNST vs. PNF tumors. The pathway as shown in aggregate score, is upregulated in MPNST vs. PNF (Wilcox test, *P* < 0.05).

We specifically compared gene expression between the MPNST and PNF cohorts. We found that *ERBB3, CNP, PMP22, PDGFB,* and *TGFBR2* were significantly downregulated in MPNST compared to PNF, while *CCNB2, TWIST1, COL6A3, PTK7, GAS1,* and *TGFB2* were significantly upregulated in MPNST (**Figure 4B**) (|fold change| > 1.2 with adjusted *P* < 0.05), consistent with previous reports [29, 30].

To validate the transcriptional programs implicated in MPNST biology, we employed gene set enrichment analysis (GSEA) on all genes differentially expressed between MPNST and PNF in our cohort. Our analysis revealed alterations in pathways within Hallmark gene sets in MPNST versus PNF (adjusted *P* < 0.01) (**Figure 4C** and **Supplementary Table 3**) (32). The three most statistically significant pathways involved upregulation of genes related to epithelial to mesenchymal transition (EMT), G2M checkpoint, and myogenesis, with volcano plots depicting the involved genes for EMT shown in **Figure 4D**. Heatmaps showing corresponding gene expression are shown in **Figure S1 A,B,C,** and differentially expressed genes in these pathways are listed in **Supplementary Tables 5, 6, 7** (adjusted *P* < 0.01 & |FC|>1.2). We also validated pathways within Reactome gene sets (33) and identified 23 Reactome pathways that were significantly different in MPNST vs. PNF (adjusted *P* < 0.01) (**Figure S2A and Supplementary Table 4**). The three most statistically significant pathways were associated with upregulation of genes involved in extracellular matrix (ECM) organization, ECM degradation, and collagen degradation, including, in particular, *MMP1*, *MMP3*, *MMP10*, and *MMP13*, members of the family of matrix metalloproteinases (34–36). Genes involved in these pathways are shown in the volcano plots in **Figure S2B,C,D** and corresponding heatmaps of gene expression are shown in **Figure S2E,F,G**. Differentially expressed genes in these pathways are listed in **Supplementary Tables 8, 9, 10** (adjusted *P* < 0.01 and |FC| > 1.2) and include multiple members of MMP and collagen gene families. These findings suggest a stronger mesenchymal phenotype in MPNST compared to pNF, and that MPNST display increased ECM and activated collagen degradation, potentially fostering tumor metastasis, as previously reported (37–39).

We also found significant activation of several pathways previously reported in the literature, which have clear links to MPNST biology including “H3K27me3-associated”, “metastasis-associated”, and “mTOR-associated” pathways in MPNST (**Supplementary Figure 3**). The single sample gene signature score for each of these gene sets was significantly higher in MPNST versus PNF (**Figure 4E** and **Supplementary Figure 3GH**, p<0.01, Wilcox test).

#### Usage Notes

All genomic datasets (both raw data and processed data) released with this article can be found on the NF Data Portal: https://bit.ly/nfbiobankdatasets. In the portal, navigating to each dataset card and clicking on the download icon in the top right will automatically add all files in the dataset for download and prompt users to complete any required data use statements to gain access and download the raw and processed data.

## Code availability

All the figures in this article are generated through reproducible R code which is available on Github (https://github.com/nf-osi/biobank-release-2). All raw data files were processed for analyses using nf-core pipelines available through https://nf-co.re/ and specific versions used in this study have been noted in the methods section.

## Acknowledgements

Funding: This publication was supported by funding from the Neurofibromatosis Therapeutic Acceleration Program (NTAP) (https://www.n-tap.org/) at the Johns Hopkins University School of Medicine, to C.A.P. and to A.C.H (Grant ID: 212027, 152009). Its contents are solely the responsibility of the authors and do not necessarily represent the official views of The Johns Hopkins University School of Medicine. Bioinformatics analyses were supported by funding from NTAP to R.J.A. and J.B (Grant ID: 229101). The funders were involved in reviewing the manuscript and providing feedback.

## Author Contributions

Authorship was determined using ICMJE recommendations.

Conceptualization: JB, ACH, JOB, CAP

Data curation: JB, YL, SCM, AC, KP, LZ

Formal Analysis: JB, YL, AJS

Funding acquisition: JOB, RJA, JB, CAP

Investigation: all authors

Methodology: JB, YL, SCM, AC, KP, LZ, AJS, ASL, AJB

Project administration: ACH, JOB, CAP

Resources: JB, YL

Software: JB, YL, AJS

Supervision: ACH, JOB, RJA, CAP

Validation: not applicable

Visualization: JB, YL, AJS

Writing: original draft: JB, YL, SCM, AJS, ACH, CAP

Writing: review & editing: all authors

## References

1. Miettinen MM, Antonescu CR, Fletcher CDM, Kim A, Lazar AJ, Quezado MM, et al. Histopathologic evaluation of atypical neurofibromatous tumors and their transformation into malignant peripheral nerve sheath tumor in patients with neurofibromatosis 1-a consensus overview. Hum Pathol. 2017;67:1–10.

2. Dehner C, Moon CI, Zhang X, Zhou Z, Miller C, Xu H, et al. Chromosome 8 gain is associated with high-grade transformation in MPNST. JCI Insight. 2021;6(6).

3. Wang J, Pollard K, Calizo A, Pratilas CA. Activation of Receptor Tyrosine Kinases Mediates Acquired Resistance to MEK Inhibition in Malignant Peripheral Nerve Sheath Tumors. Cancer Res. 2021;81(3):747–62.

4. Larsson AT, Bhatia H, Calizo A, Pollard K, Zhang X, Conniff E, et al. Ex vivo to in vivo model of malignant peripheral nerve sheath tumors for precision oncology. Neuro Oncol. 2023;25(11):2044–57.

5. Pollard K, Banerjee J, Doan X, Wang J, Guo X, Allaway R, et al. A clinically and genomically annotated nerve sheath tumor biospecimen repository. Sci Data. 2020;7(1):184.

6. Legius E, Messiaen L, Wolkenstein P, Pancza P, Avery RA, Berman Y, et al. Revised diagnostic criteria for neurofibromatosis type 1 and Legius syndrome: an international consensus recommendation. Genet Med. 2021;23(8):1506–13.

7. Li H. Aligning sequence reads, clone sequences and assembly contigs with BWA-MEM 2013 [

8. McKenna A, Hanna M, Banks E, Sivachenko A, Cibulskis K, Kernytsky A, et al. The Genome Analysis Toolkit: a MapReduce framework for analyzing next-generation DNA sequencing data. Genome Res. 2010;20(9):1297–303.

9. DePristo MA, Banks E, Poplin R, Garimella KV, Maguire JR, Hartl C, et al. A framework for variation discovery and genotyping using next-generation DNA sequencing data. Nat Genet. 2011;43(5):491–8.

10. Van der Auwera GA, Carneiro MO, Hartl C, Poplin R, Del Angel G, Levy-Moonshine A, et al. From FastQ data to high confidence variant calls: the Genome Analysis Toolkit best practices pipeline. Curr Protoc Bioinformatics. 2013;43(1110):11 0 1–0 33.

11. Kim S, Scheffler K, Halpern AL, Bekritsky MA, Noh E, Kallberg M, et al. Strelka2: fast and accurate calling of germline and somatic variants. Nat Methods. 2018;15(8):591–4.

12. Cibulskis K, Lawrence MS, Carter SL, Sivachenko A, Jaffe D, Sougnez C, et al. Sensitive detection of somatic point mutations in impure and heterogeneous cancer samples. Nat Biotechnol. 2013;31(3):213–9.

13. Kandoth C, Gao JJ, Qwang M, Mattioni M, Struck A, Boursin Y, et al. mskcc/vcf2maf: vcf2maf v1.6.16 2018 [Available from: doi:10.5281/zenodo.593251.

14. Garcia M, Juhos S, Larsson M, Olason PI, Martin M, Eisfeldt J, et al. Sarek: A portable workflow for whole-genome sequencing analysis of germline and somatic variants. F1000Res. 2020;9:63.

15. Kuhn RM, Haussler D, Kent WJ. The UCSC genome browser and associated tools. Brief Bioinform. 2013;14(2):144–61.

16. Fang LT, Afshar PT, Chhibber A, Mohiyuddin M, Fan Y, Mu JC, et al. An ensemble approach to accurately detect somatic mutations using SomaticSeq. Genome Biol. 2015;16(1):197.

17. Patel HE, P., Peltzer, A. Botvinnik, O., Manning, J., Sturm, G., Garcia, M. Moreno, D. Vemuri, P., nf-core bot, Pantano, L., Binzer-Panchal, M., Zepper, M., Syme, R., Kelly, G., Hanssen, F., rfenouil, Espinosa-Carrasco, J., marchoeppner, Miller, E., Zhou, P., Guinchard, S., Gabarnet, G., Mertes, C., Straub, D., DiTommaso, P. nf-core/rnaseq: nf-core/rnaseq v3.13.2 - Cobalt Colt 2023 [updated November 21, 2023. Version 3.13.2:[

18. Patro R, Duggal G, Love MI, Irizarry RA, Kingsford C. Salmon provides fast and bias-aware quantification of transcript expression. Nat Methods. 2017;14(4):417–9.

19. Leek JT, Johnson WE, Parker HS, Jaffe AE, Storey JD. The sva package for removing batch effects and other unwanted variation in high-throughput experiments. Bioinformatics. 2012;28(6):882–3.

20. McInnes L, Healy J, Melville J. UMAP: Uniform Manifold Approximation and Projection for Dimension Reduction: arXiv; 2018 [

21. Foroutan M, Bhuva DD, Lyu R, Horan K, Cursons J, Davis MJ. Single sample scoring of molecular phenotypes. BMC Bioinformatics. 2018;19(1):404.

22. Pedersen BS, Bhetariya PJ, Brown J, Kravitz SN, Marth G, Jensen RL, et al. Somalier: rapid relatedness estimation for cancer and germline studies using efficient genome sketches. Genome Med. 2020;12(1):62.

23. Zhang M, Wang Y, Jones S, Sausen M, McMahon K, Sharma R, et al. Somatic mutations of SUZ12 in malignant peripheral nerve sheath tumors. Nat Genet. 2014;46(11):1170–2.

24. Lee W, Teckie S, Wiesner T, Ran L, Prieto Granada CN, Lin M, et al. PRC2 is recurrently inactivated through EED or SUZ12 loss in malignant peripheral nerve sheath tumors. Nat Genet. 2014;46(11):1227–32.

25. De Raedt T, Beert E, Pasmant E, Luscan A, Brems H, Ortonne N, et al. PRC2 loss amplifies Ras-driven transcription and confers sensitivity to BRD4-based therapies. Nature. 2014;514(7521):247-51.

26. Sohier P, Luscan A, Lloyd A, Ashelford K, Laurendeau I, Briand-Suleau A, et al. Confirmation of mutation landscape of NF1-associated malignant peripheral nerve sheath tumors. Genes Chromosomes Cancer. 2017;56(5):421–6.

27. Cortes-Ciriano I, Steele CD, Piculell K, Al-Ibraheemi A, Eulo V, Bui MM, et al. Genomic Patterns of Malignant Peripheral Nerve Sheath Tumor (MPNST) Evolution Correlate with Clinical Outcome and Are Detectable in Cell-Free DNA. Cancer Discov. 2023;13(3):654–71.

28. McLaren W, Gil L, Hunt SE, Riat HS, Ritchie GR, Thormann A, et al. The Ensembl Variant Effect Predictor. Genome Biol. 2016;17(1):122.

29. Mo J, Moye SL, McKay RM, Le LQ. Neurofibromin and suppression of tumorigenesis: beyond the GAP. Oncogene. 2022;41(9):1235–51.

30. Bottillo I, Ahlquist T, Brekke H, Danielsen SA, van den Berg E, Mertens F, et al. Germline and somatic NF1 mutations in sporadic and NF1-associated malignant peripheral nerve sheath tumours. J Pathol. 2009;217(5):693–701.

31. Upadhyaya M, Spurlock G, Monem B, Thomas N, Friedrich RE, Kluwe L, et al. Germline and somatic NF1 gene mutations in plexiform neurofibromas. Hum Mutat. 2008;29(8):E103–11.

32. Liberzon A, Birger C, Thorvaldsdottir H, Ghandi M, Mesirov JP, Tamayo P. The Molecular Signatures Database (MSigDB) hallmark gene set collection. Cell Syst. 2015;1(6):417–25.

33. Gillespie M, Jassal B, Stephan R, Milacic M, Rothfels K, Senff-Ribeiro A, et al. The reactome pathway knowledgebase 2022. Nucleic Acids Res. 2022;50(D1):D687–D92.

34. Shapiro SD. Matrix metalloproteinase degradation of extracellular matrix: biological consequences. Curr Opin Cell Biol. 1998;10(5):602–8.

35. Frantz C, Stewart KM, Weaver VM. The extracellular matrix at a glance. J Cell Sci. 2010;123(Pt 24):4195–200.

36. Lu P, Takai K, Weaver VM, Werb Z. Extracellular matrix degradation and remodeling in development and disease. Cold Spring Harb Perspect Biol. 2011;3(12).

37. Brockman QR, Scherer A, McGivney GR, Gutierrez WR, Voigt AP, Isaacson AL, et al. PRC2 loss drives MPNST metastasis and matrix remodeling. JCI Insight. 2022;7(20).

38. Thomson CS, Pundavela J, Perrino MR, Coover RA, Choi K, Chaney KE, et al. WNT5A inhibition alters the malignant peripheral nerve sheath tumor microenvironment and enhances tumor growth. Oncogene. 2021;40(24):4229–41.

39. Errico A, Stocco A, Riccardi VM, Gambalunga A, Bassetto F, Grigatti M, et al. Neurofibromin Deficiency and Extracellular Matrix Cooperate to Increase Transforming Potential through FAK-Dependent Signaling. Cancers (Basel). 2021;13(10).

